# Predicting cellular functions of human phosphosites

**DOI:** 10.64898/2026.05.22.727149

**Authors:** Julian van Gerwen, Danielle Gallagher, Irene Sala, Jürgen Jänes, Kosuke Ogata, Jacob E. Corn, Pedro Beltrao

## Abstract

Protein phosphorylation is a regulatory switch that controls essentially all cellular processes, including metabolism, cell cycle progression, and DNA repair. Furthermore, by modulating specific regions and activities of a protein, phosphorylation sites can independently coordinate the protein’s involvement in distinct cellular processes. Despite the importance of this regulatory mechanism, the majority of human phosphosites have not been functionally characterized. Here we have developed biologically informed predictors of phosphosite cellular function by integrating protein functional annotation, recognition by upstream kinases, and regulation across phosphoproteomics perturbations. To enable this we harmonized public phosphoproteomics data into an atlas of nearly 2,000 perturbations and more than 16 million quantifications, providing a powerful resource for studying phosphosite regulation. We applied our predictors to functionally characterise thousands of understudied phosphosites, revealing many phosphosite functions that could not be inferred from protein functional annotation, and functionally divergent phosphosites within the same protein. Using cancer multi-omics data we characterized connections between these phosphosites and other molecular layers. Finally, we illuminated novel signaling axes within the DNA repair pathway, and demonstrated that phosphosites on PPM1G and KHSRP modulate cell survival under DNA damage. Our work represents a powerful resource for dissecting the cellular consequences of protein phosphorylation.

## Introduction

Reversible protein phosphorylation allows cells to rapidly sense and respond to changes in their environment. Environmental signals regulate cascades of protein kinases and phosphatases, which catalyse the addition and removal of phosphates on specific serine, threonine, or tyrosine residues in eukaryotic proteins. Phosphorylation alters a protein’s molecular function through several avenues, including tuning enzymatic activity, promoting or disrupting protein-protein interactions, and regulating subcellular localisation^1^. The coordinated modulation of specific proteins regulates cellular pathways, such as cell growth and cell cycle progression, resulting in adaptation to the environmental change^1^. Protein phosphorylation regulates essentially every cellular process and is dysregulated in myriad diseases including cancer^2^, making it a critical target of biomolecular research.

Over the last two decades, our understanding of protein phosphorylation has been greatly advanced by phosphoproteomics - the unbiased identification and quantification of phosphorylation by mass spectrometry. Phosphoproteomics has uncovered more than 100,000 unique human phosphosites, distributed through the majority of the proteome^3^. However, the rapid discovery afforded by phosphoproteomics has outpaced functional characterisation. Only approximately 3% of human phosphosites have been assigned an experimentally determined function, which is traditionally achieved by low-throughput site mutagenesis coupled to biomolecular characterisation, and only 3% have been assigned an upstream kinase^2^. This constitutes a major barrier to understanding how the phosphoproteome controls cell biology.

Studying phosphosite function has been complicated by the possibility that many phosphosites may be functionally irrelevant. Indeed, protein phosphorylation is energetically cheap and phosphosites generally evolve rapidly, suggesting that many are non-functional evolutionary noise^4^. Encouragingly, computational tools have begun to predict which phosphosites are most critical for the cell^3,5–7^, and experimental approaches employing high-throughput site mutagenesis or functional proteomics are corroborating these predictions at growing scales^8–13^. Now that we can prioritise phosphosites potentially important to the cell, we next need approaches to determine which cellular processes they regulate, such as cell cycle progression, cell growth, and DNA repair. Phenotypic screens that assay phosphosite mutation across diverse conditions provide rich information suitable for inferring phosphosite function^8^. Alternatively, protein and phosphosite association networks have been employed to predict phosphosite function by guilt-by-association^14^, however the utility of additional information and approaches to computationally infer phosphosite function has not been assessed. Overall, the global prediction of phosphosite cellular functions remains underexplored.

Here, we have addressed the challenge of predicting the cellular functions of understudied phosphosites. Examining experimentally determined relationships between human phosphosites and the cellular processes they regulate, we uncovered associations from processes to protein-level ontology and upstream kinase specificity. Hypothesising that the regulation of phosphosites under specific conditions would be indicative of their function, we compiled public phosphoproteomics data comprising nearly 2,000 control-matched perturbations, quantifying 189,576 human phosphosites and over 16,000,000 phosphosite changes. We then leveraged these features to construct machine learning models of phosphosite function and predicted cellular processes for thousands of understudied phosphosites. We explored systems-level consequences of these phosphosites in cancer multi-omics data, illuminated canonical and non-canonical signaling axes within the DNA repair pathway, and validated the contribution of phosphosites on PPM1G and KHSRP to cell survival under DNA damage. Our work offers insights into the phosphoregulation of cellular processes, and provides a powerful resource for functional analysis of protein phosphorylation.

## Results

### Phosphosite cellular function associates with protein annotation and kinase specificity

Site-specific phosphorylation of serine, threonine, and tyrosine residues regulates essentially every cellular process, such as cell cycle progression, apoptosis, and DNA repair. We first set out to determine the principles connecting phosphosites to the specific cellular processes they regulate, in order to inform the design of machine-learning predictors of phosphosite cellular function. The PhosphositePlus repository curates regulatory phosphosites, which have been experimentally determined to regulate a diverse set of cellular processes (Fig. 1a^15^). We focused on the 12 processes with the most annotated regulatory phosphosites, comprising 6,334 total phosphosites in humans (Fig. 1a-b, Table S1). More than half of these phosphosites regulate only one cellular process, suggesting these annotations can elucidate the unique regulatory profile of each process (Fig. 1b).

**Figure 1:**
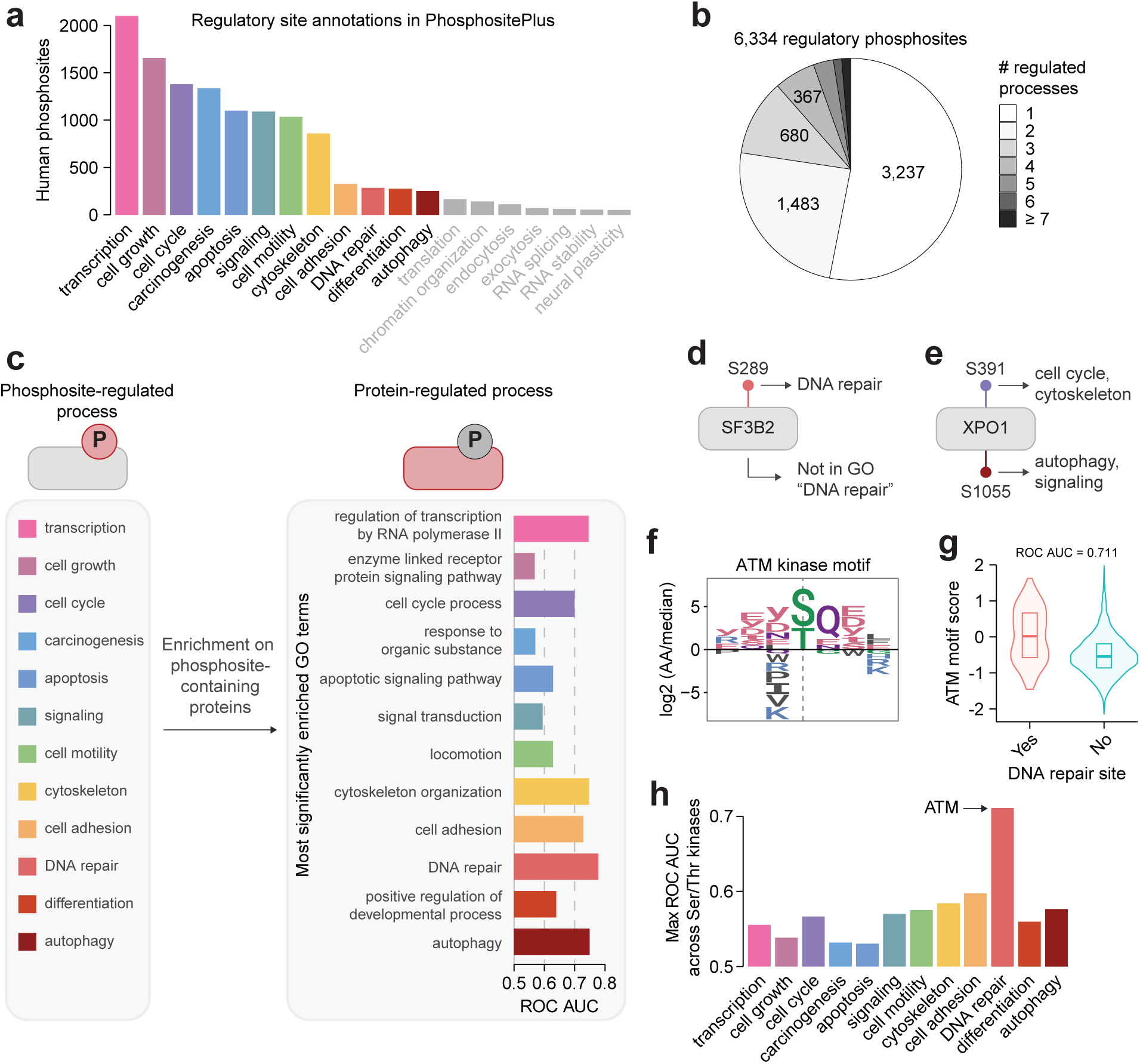
Phosphosite cellular function associates with protein annotation and kinase specificity. **A)** The number of human phosphosites annotated to regulate cellular processes in PhosphositePlus^15^. **B)** The number of cellular processes annotated to each unique human phosphosite. **C)** The most significantly enriched GO biological process terms among proteins containing process-regulating phosphosites. ROC AUCs were calculated for using GO terms as predictors of phosphosite function. **D-E)** Two proteins with regulatory phosphosites that could not be predicted with GO terms. The processes regulated by each phosphosite are shown. **F)** The sequence motif of the ATM kinase from Johnson et al.^16^. **G)** ATM motif scores for Ser/Thr phosphosites that regulate DNA repair or other cellular processes. The ROC AUC was calculated for predicting DNA repair sites with the motif score. **H)** Motifs for 303 Ser/Thr kinases^16^ were used to predict phosphosites regulating each process. The maximum ROC AUC across kinases is shown.

The cellular function of a phosphosite should be strongly dictated by that of the protein it modifies. For instance, proteins containing phosphosites that regulate transcription were enriched in concordant gene ontology (GO) biological process terms, such as “regulation of transcription by RNA polymerase II” and “transcription, DNA templated” (Fig. 1c, Table S2). In general we observed similar concordant enrichment for phosphoproteins regulating the other 11 cellular processes (Fig. 1c). Despite this, when using the most significantly enriched GO term from each process to predict its regulatory phosphosites, we achieved only moderate accuracy (Fig. 1c). Indeed, this approach fails on understudied proteins that lack the requisite GO annotation - such as the splicing factor SF3B2, which contains a DNA repair-regulating phosphosite but lacks GO “DNA repair” annotation (Fig. 1d) - or when multiple phosphosites within a protein regulate distinct processes - such as the nuclear export protein XPO1, which contains one phosphosite regulating the cell cycle and cytoskeletal reorganisation, and another phosphosite regulating autophagy and signaling (Fig. 1e).

Beyond protein-level information, the regulation of a phosphosite by upstream kinases may further inform on its function, as kinases can coordinately modulate cellular pathways during signal transduction. Since the *in vivo* upstream kinases of most phosphosites are unknown^2^, we took advantage of substrate sequence specificities, which were recently resolved for nearly all human kinases and can be applied to any phosphosite as an approximation of substrate recognition^16,17^. For example, ATM, ATR, and DNA-PK - key coordinators of the DNA damage response - uniquely target Ser/Thr sites followed by glutamine, and this motif matched DNA repair-regulating phosphosites better than other regulatory phosphosites (Fig. 1f-g). Beyond DNA repair, Ser/Thr kinase specificities could only weakly predict phosphosite cellular function (Fig. 1h). However, in general the most predictive kinases were still enriched in known regulators of the corresponding cellular process, such as CDK kinases for the cell cycle, and PAK and ROCK kinases for the cytoskeleton (Fig. S1a). Overall, phosphosite cellular function cannot be predicted entirely from protein annotation, and is generally weakly associated with kinase sequence specificity.

### An atlas of perturbation phosphoproteomics

Since protein phosphorylation transduces environmental perturbations to cellular adaptations, we reasoned that the dynamic regulation of phosphosites could illuminate their cellular function. While phosphosite regulation is routinely measured by mass spectrometry-based phosphoproteomics, most studies only examine a few distinct conditions. This means that we lack a comprehensive map of the phosphoproteome across diverse cellular perturbations. To address this, we collated published phosphoproteomics experiments measuring control-matched perturbations from three sources: the qPTM database^18^, our previous compilation of perturbation experiments^19^, and an updated curation effort whereby we collected 27 major phosphoproteomics studies published since 2016^20–46^ (Fig. 2a, see Methods). After data harmonisation, our phosphoproteomics atlas comprised 1,937 perturbations, 189,576 human phosphosites, and 16,157,226 quantitative fold-change values (Fig. 2b-c, Table S3).

**Figure 2:**
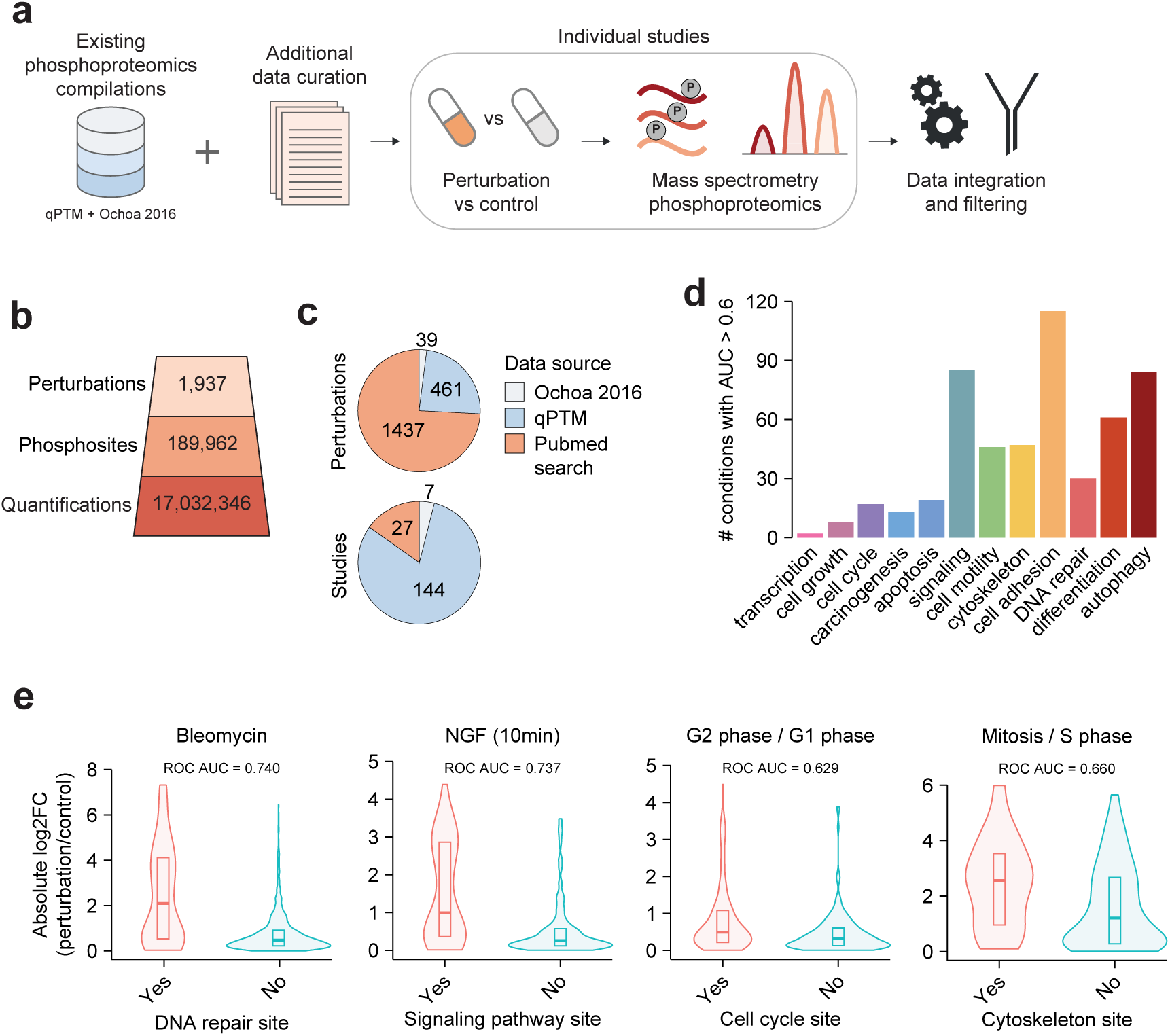
An atlas of phosphoproteomics perturbations. **A)** Workflow for compiling phosphoproteomics experiments of control-matched perturbations into a phosphoproteomics atlas. **B)** Number of perturbations, quantified phosphosites, and perturbation/control fold-change values (“Quantifications”) in the phosphoproteomics atlas. **C)** The source of perturbations and studies in the phosphoproteomics atlas. Perturbations and studies present in both Ochoa 2016 and qPTM are labelled as qPTM. **D)** The absolute log2-fold change under each perturbation was used to predict phosphosites regulating each process.Conditions with fewer than 26 phosphosites quantified in the positive or negative class were removed. **E)** Example perturbations that predicted phosphosites regulating distinct cellular processes.

Our atlas contained many perturbations that regulated specific cellular processes (Fig. 2d). These included expected relationships, for example, the DNA damage agent bleomycin preferentially regulated DNA repair phosphosites; the growth factor NGF regulated sites involved in signaling pathway regulation; quantitative comparison of cells in G2 vs G1 phase enriched for cell cycle-regulating phosphosites; and comparison of cells in mitosis vs S phase enriched for sites involved in cytoskeletal dynamics (Fig. 2e). Based on this, we expect our atlas can also uncover unknown connections between specific cellular perturbations and the biological processes they regulate. For instance, the cough medication noscapine caused changes in phosphorylation that were enriched for cell cycle phosphosites, which is supported by evidence that noscapine induces mitotic arrest^47^ (Fig. S2a). Furthermore, changes induced by the tyrosine phosphatase inhibitor pervanadate were enriched in cell adhesion phosphosites, consistent with experiments where pervanadate induced T-cell adhesion^48^ (Fig. S2b). In all, our atlas provides a phosphoproteomic map from biological perturbations to cellular biology.

### Predicting phosphosite cellular function

We next sought to integrate phosphoproteomic regulation, kinase sequence specificity, and protein annotation into unified predictors of phosphosite cellular function. To train a model predicting the phosphosites regulating a given cellular process (referred to here as “process X”), we would ideally compare a set of sites known to regulate process X with sites that do not regulate process X. However, there is very little definitive evidence proving a site does not regulate a given process. Instead, as a negative set we use all phosphosites annotated to regulate any process apart from process X, which is justified by the assumptions that these sites are less likely to regulate process X given they have already been implicated in another process, and that phosphosites regulating process X will represent only a fraction of the phosphoproteome and hence would be rare in this group.

Specifically, we first reserved 20% of PhosphositePlus regulatory phosphosites as a test set and performed feature selection for each cellular process on the remaining sites (Fig. 3a). To combat data sparseness we focused on Ser/Thr phosphosites quantified in at least three of the selected phosphoproteomics features, and imputed the remaining missing values using the mean of each feature. For each cellular process we then trained an independent binary classifier to discriminate phosphosites regulating a given process from those regulating other processes, which allows us to predict phosphosites regulating multiple processes. We repeated this procedure over 50 random train-test splits for robust estimates of model accuracy.

**Figure 3:**
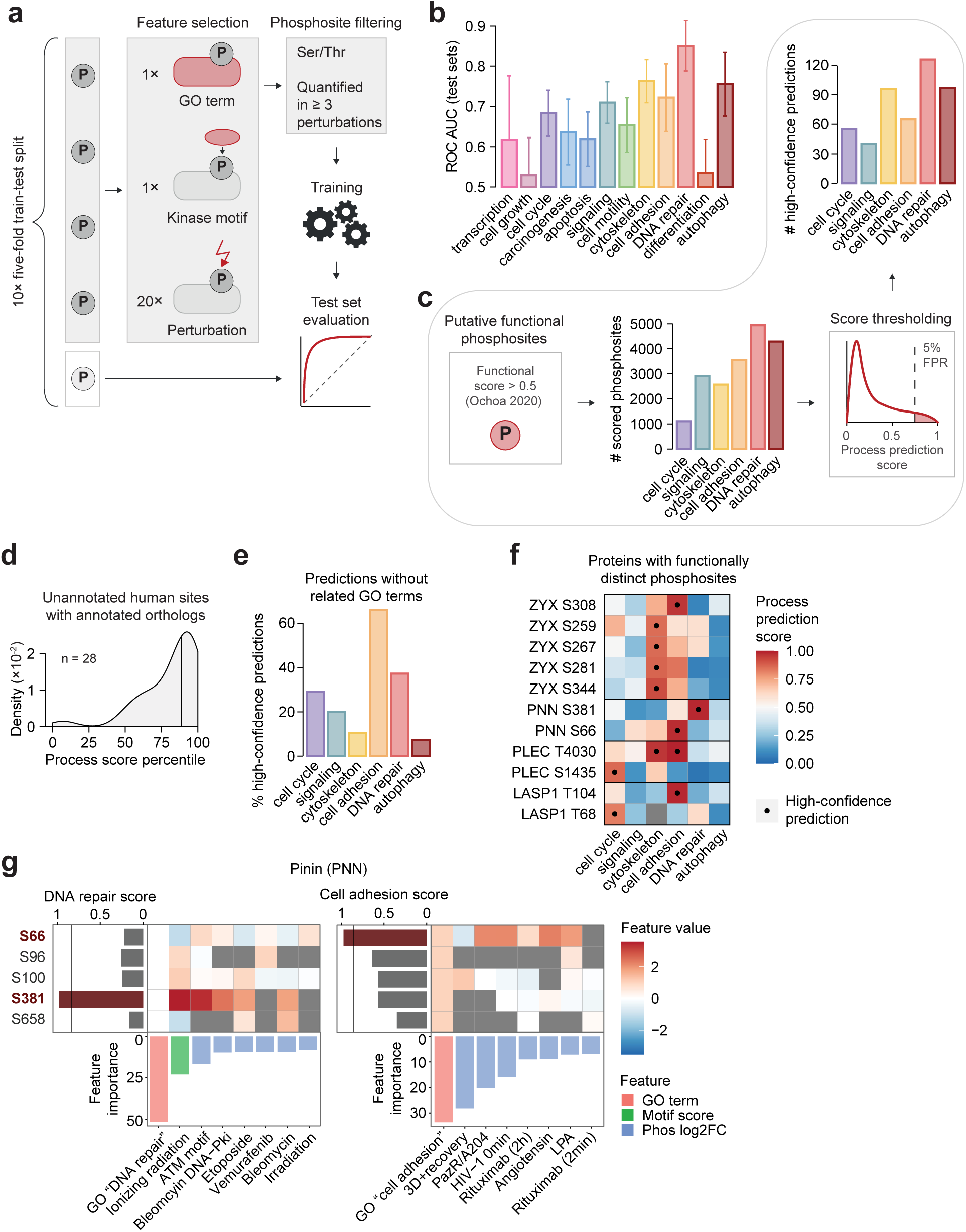
Predicting phosphosite cellular function. **A)** Workflow for training predictors of phosphosite cellular function. **B)** Performance of cellular function predictors across 50 randomly selected test sets (80:20 train:test split). Error bars indicate standard deviation. **C)** Predictors were applied to putative regulatory phosphosites (Ochoa 2020 functional score > 0.5^3^), resulting in thousands of scored phosphosites for each process. Scores were thresholded at 5% FPR based on test-set performance, to obtain high-confidence predictions. **D)** Unannotated human phosphosites scored highly for cellular processes regulated by orthologous sites in other species. The percentiles of phosphosites within each process score distribution are shown in order to visualise all processes together. 33 unique phosphosites are shown. **E)** The percentage of high-confidence predictions that were on proteins not annotated to any GO terms used by the corresponding predictors. **F)** Proteins with multiple high-confidence phosphosites predicted for different cellular processes. A score difference between processes of > 0.25 was required. **G)** DNA repair and cell adhesion prediction scores for phosphosites in the protein Pinin. Raw values for the eight most important features are shown. High-confidence predictions are highlighted in dark red (S381 predicted to regulate DNA repair, S66 predicted to regulate cell adhesion).

We achieved similar performance using logistic regression and gradient-boosting machine models, hence we selected logistic regression due to its interpretability (Fig. S3a). Cellular processes differed in their predictability, ranging from DNA repair with a mean test-set ROC AUC of 0.85, to cell growth with 0.52 (Fig. 3b). In addition to cell growth, the other poorly performing processes such as differentiation and carcinogenesis were typically broader in their biological scope, and hence may be harder to predict with our approach. For further analysis we selected the top 6 processes, which featured mean ROC AUCs of approximately 0.7 or higher. Importantly, models trained on all feature modalities generally outperformed models trained just on GO biological process terms, highlighting the importance of integrating phosphosite-level information in addition to protein function (Fig. S3a).

We next employed these models to characterise the unstudied phosphoproteome. Since many unstudied phosphosites may serve no major regulatory role, we focused on putative regulatory phosphosites using our recently published functional score (score > 0.5^3^, Fig. 3c). Applying the six most accurate cellular-process models, we scored thousands of these phosphosites for their propensity to regulate each process, noting that each model could not be applied to exactly the same set of sites due to different phosphoproteomics conditions being employed (Fig. 3c, Table S4). Finally, to ensure the reliability of downstream analysis, we obtained high-confidence predictions by setting a stringent score cutoff (FPR < 5%), comprising 41-117 phosphosites per cellular process (Fig. 3c, Table S4).

We identified 28 unannotated human phosphosites where the orthologous phosphosite in another species was functionally annotated. Under the assumption that the functions of these phosphosites are conserved in humans, they can serve as an orthogonal validation set, since they were not used in training. The 28 human phosphosites generally scored highly for the cellular process regulated by their ortholog, confirming the accuracy of our predictions (Fig. 3d). For example, DNA repair is regulated by the mouse phosphosite Cops3 S410 and predicted to be regulated by the unannotated human ortholog COPS3 S410 (score = 0.878). In all, we have trained accurate models of phosphosite cellular function and applied them to the unstudied phosphoproteome.

### Interpreting predictions of phosphosite cellular function

Interrogating the unbiased feature selection of our models provides insight into their functional predictions. Encouragingly, the models incorporated protein-level GO terms that aligned with the predicted cellular process, such as “DNA repair” and “cell cycle process” for the DNA repair and cell cycle models respectively (Table S5). Despite this, a substantial fraction of high-confidence function predictions were for phosphosites on proteins lacking the relevant GO annotation, corresponding to 40% and 30% of DNA repair and cell cycle predictions, for example (Fig. 3e). Hence, kinase-motif and phosphoproteomics features provide novelty by identifying phosphosite cellular functions on non-canonical proteins not yet implicated in the corresponding cellular process.

Another phenomenon that cannot be predicted from protein-level function annotation is when multiple phosphosites control distinct functions of a single protein. Excitingly, we were able to predict four of this behaviour (Fig. 3f). For example, we predicted that the phosphosites S66 and S381 in the protein Pinin (PNN) regulate cell adhesion and DNA repair respectively, which aligns with the well-established role of Pinin in regulating cell adhesion through E-cadherin transcription^49^, and evidence that PNN participates in DNA repair in *C. elegans*^50^ (Fig. 3f). Additionally, we predicted Plectin (PLEC) S1435 to regulate cell cycle and T4030 to regulate cell adhesion and the cytoskeleton, which agrees with the roles of Plectin in cytoskeletal rearrangements during mitosis^51^ and anchoring focal adhesion to intermediate filaments^52^ (Fig. 3f).

An additional benefit of our biologically informed features is that they lend interpretability to our predictions. For instance, while Pinin lacks GO “DNA repair” annotation - the most heavily weighted feature for DNA repair prediction - S381 strongly matches the ATM kinase motif and is up-regulated in heavily weighted phosphoproteomics conditions, which mostly comprise treatment with DNA damage agents (Fig. 3g). This provides interpretable and compelling evidence that S381 is involved in the DNA damage response. To understand why S66 is predicted to regulate cell adhesion, one can observe that Pinin is annotated to “cell adhesion” in GO, and S66 is additionally up-regulated in some of the most heavily weighted phosphoproteomics conditions, some of which have demonstrated links to cell adhesion (e.g. LPA^53^) (Fig. 3g). Overall, our predictions present a wealth of interpretable information beyond existing annotations, and reveal the regulatory nuances of phosphorylation in coordinating multi-functional proteins.

### Contextualising predictions in cancer multi-omics data

Next, in order to understand how our predicted phosphosite functions interact with other molecular layers such as gene expression or mutations, we turned to CPTAC, a collection of multi-omics data featuring over 1,000 tumour phosphoproteomes from 10 tumour types^54^, which was not used in our model training (Fig. 4a). Interestingly, while these data contained less than half of the phosphosites annotated with regulatory functions in PhosphositePlus, nearly all phosphosites that we assigned high-confidence function predictions were quantified, likely due to the incorporation of phosphoproteomics data as model features (Fig. 4b). Hence, our predictions have high potential for facilitating the functional analysis of phosphoproteomics studies.

**Figure 4:**
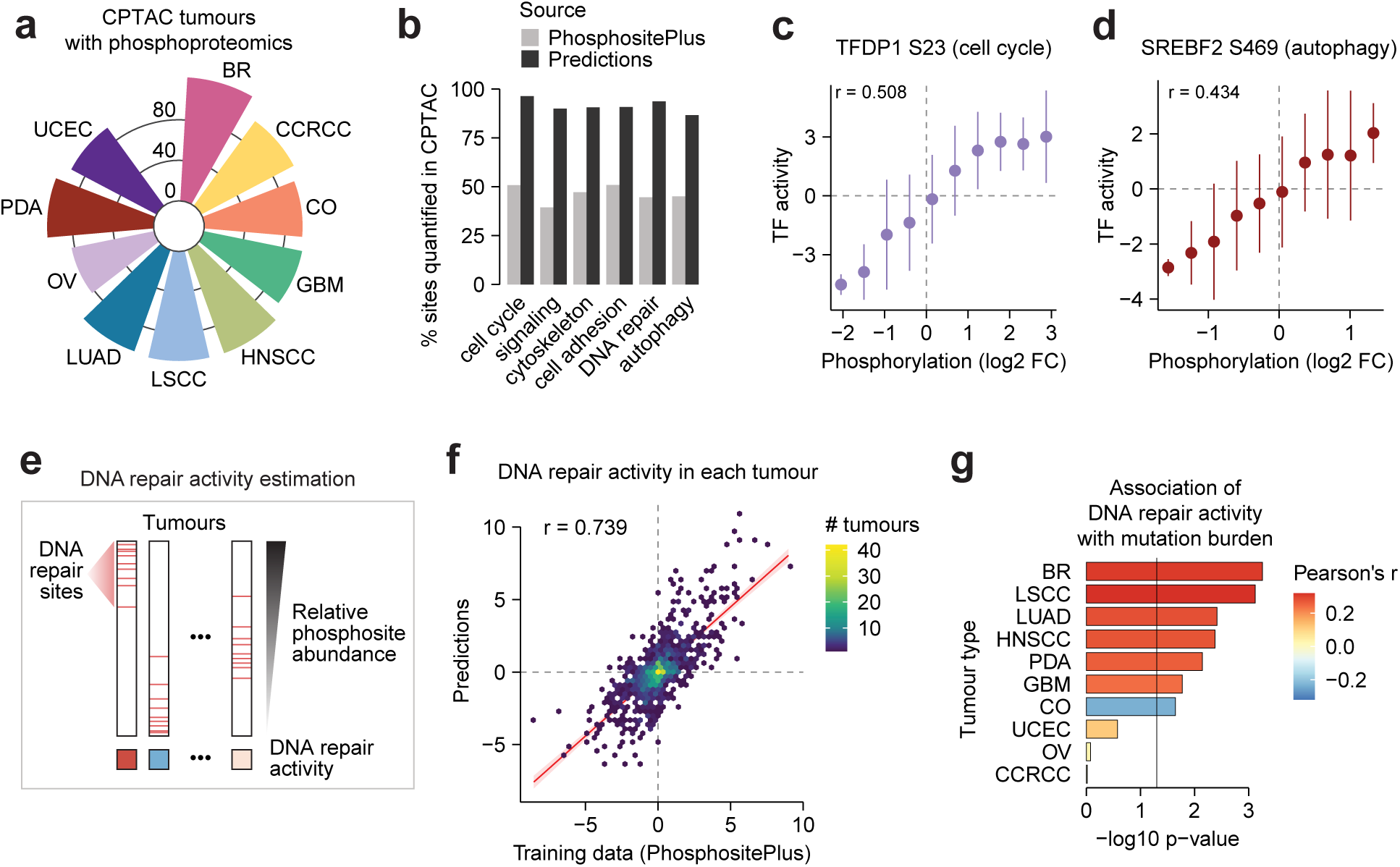
Contextualising phosphosite function predictions in cancer multi-omics data. **A)** The number of unique tumours of each tumour type measured by phosphoproteomics in CPTAC (BR: breast cancer, CCRCC: clear cell renal cell carcinoma, CO: colon cancer, GBM: glioblastoma multiforme, HNSCC: head and neck squamous cell carcinoma, LSCC: lung squamous cell carcinoma, LUAD: lung adenocarcinoma, OV: ovarian cancer, PDA: pancreatic ductal adenocarcinoma, UCEC: uterine corpus endometrial carcinoma). **B)** The percentage of PhosphositePlus-annotated sites or high-confidence predictions quantified in CPTAC phosphoproteomics data. **C-D)** High-confidence phosphosites on transcription factors, whose abundance was associated with estimated transcription factor activity by Pearson’s correlation. Points represent mean transcription factor activity in bins of phosphosite abundance and error bars represent standard deviation. **E)** Framework for estimating DNA repair signaling activity in individual tumours using relative abundance of DNA repair phosphosites. **F)** Pearson’s correlation between DNA repair signaling activity (signed log10 p-values) estimated using PhosphositePlus DNA repair regulatory sites or high-confidence DNA repair predictions. The red line indicates linear regression and shading indicates 95% confidence intervals. **G)** DNA repair activity was estimated by combining PhosphositePlus and high-confidence phosphosites and was correlated with somatic mutation burden within each tumour type by Pearson’s correlation. Vertical line indicates p = 0.05.

We next leveraged CPTAC transcriptomics data to ask whether our phosphosites modulate transcriptional programs. By estimating transcription factor activity from target transcript abundance (see Methods), we found two transcription factors with phosphosites that correlated with their activity and for which we could nominate cellular functions: TFDP1 S23, predicted to regulate cell cycle (r = 0.508, p = 2.7×10^−63^ for correlation with transcription factor activity), and SREBF2 S469, predicted to regulate autophagy (r = 0.434, p = 1.1×10^−16^, Fig. 4c-d). Importantly, activity correlated more strongly with phosphorylation than with abundance of the proteins themselves, suggesting these phosphosites may directly activate their host transcription factors (Fig. S4a). Machine learning model features revealed that TFDP1 S23 matches the CDK kinase motif and is up-regulated in late S and G2 phase, while SREBF2 is up-regulated following 12 hours of mitophagy induction with CCCP, providing further contextual evidence for the cellular roles of these phosphosites (Fig. S4b-d).

Given that DNA repair is frequently dysregulated in cancer^55^, we next estimated the overall activation of DNA repair signaling within each tumour with a rank-based enrichment test (see Methods), using either our high-confidence DNA repair-predicted phosphosites (excluding training data phosphosites) or the PhosphositePlus annotations on which they were trained (Fig. 4e). Despite the fact that CPTAC data was not used in our prediction, activity estimates from these two sources were highly correlated, validating that our predictions emulate their training data in an independent biological setting (r = 0.739, p = 1.4×10^−177^, Fig. 4f). Furthermore, DNA repair signaling activity was positively associated with somatic mutation burden in most tumour types, consistent with a plausible model whereby higher frequency of mutational events leads to heightened activation of DNA repair signaling (Fig. 4g). In all, cancer multi-omics data allowed us to interpret mechanistic consequences of our predicted phosphosite functions and further validate the biological consistency of DNA repair predictions.

### An expanded map of DNA repair signaling

We next undertook a deeper investigation into our DNA repair phosphosite function predictions. To understand the consequences when these phosphosites are dysregulated, we interrogated the ClinVar database of human disease-linked mutations^57^, identifying 255 missense mutations that disrupted phosphosites scored by our DNA repair predictor, mostly classified at the lower confidence level of “uncertain pathogenicity”. When considering mutations disrupting phosphosites with high DNA repair prediction, the most significantly enriched mutation-associated disease was “Hereditary cancer-predisposing syndrome”, which is expected given the role of DNA repair dysregulation in cancer^55^ (Fig. 5a, Table S6). The highest-scoring cancer-linked phosphosites included S397 and S738 on NBN and S20 on RAD51C, two canonical mediators of homologous recombination, highlighting these phosphosites as putative focal points linking DNA repair dysregulation to cancer progression (Fig. 5a).

**Figure 5:**
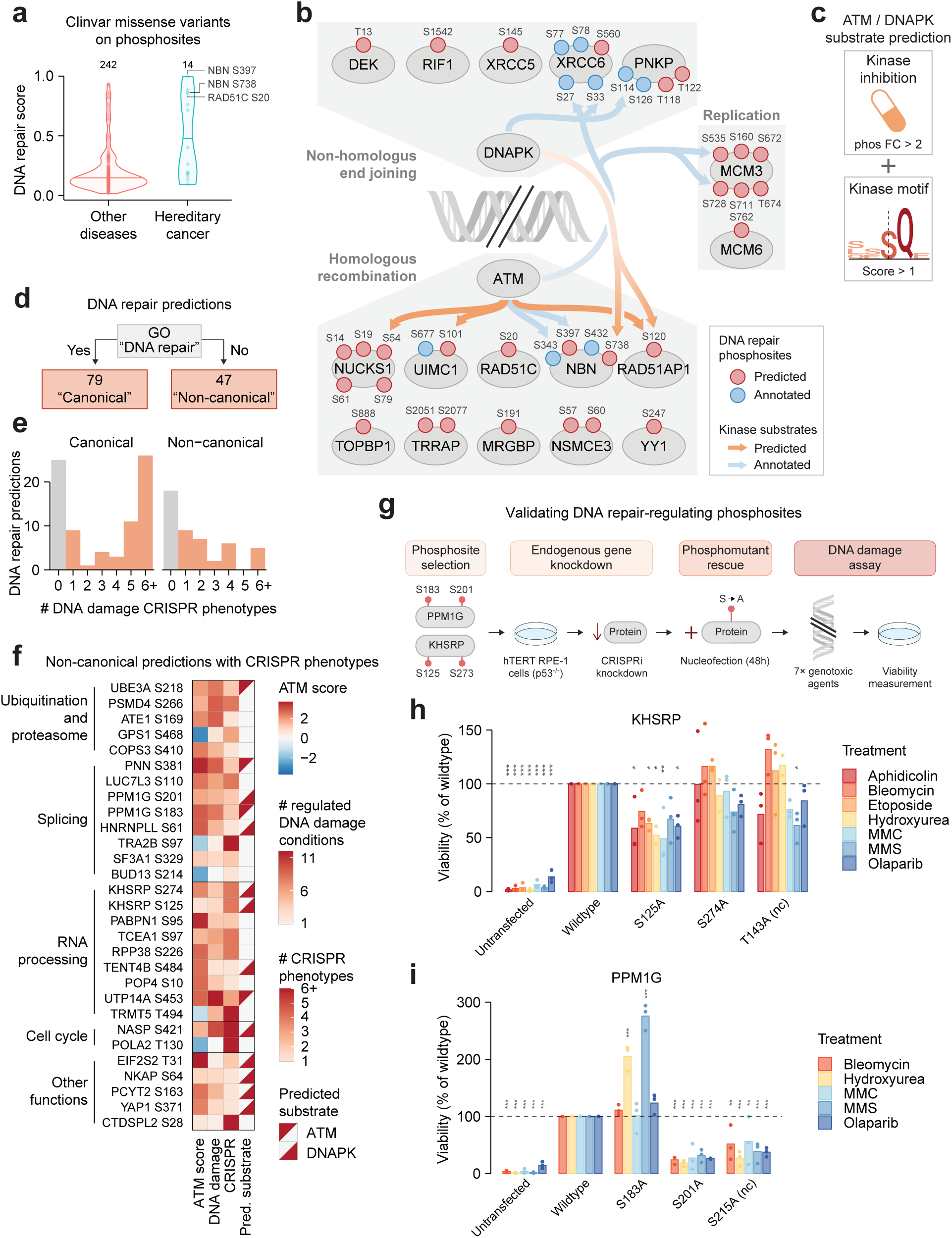
An expanded map of DNA repair signaling. **A)** DNA repair prediction scores for phosphosites that colocalized with ClinVar missense mutations. Mutations are separated by those annotated to hereditary cancer syndromes or other diseases. **B)** Proteins with high-confidence DNA repair phosphosites that participate in non-homologous end joining, homologous recombination, and DNA replication (see Methods for protein curation). High-confidence DNA repair phosphosites are colored red and PhosphositePlus DNA repair phosphosites are colored blue. Kinase-substrate relationships curated in PhosphositePlus are shown in blue and predicted relationships are shown in red. **C)** Framework for predicting DNA repair kinase substrates using sequence motifs and phosphoproteomics inhibitor data. **D)** The number of “canonical” and “non-canonical” DNA repair predictions, based on “DNA repair” GO annotation. **E)** The number of gene-level CRISPR phenotypes under DNA damage and related conditions, curated in DDRcs^61^ (FDR < 0.1). **F)** Non-canonical DNA repair predictions with DNA damage CRISPR phenotypes, grouped by protein function. ATM motif scores are shown, as well as the number of DNA damage conditions in our phosphoproteomics atlas with > 2-fold up/down-regulation, the number of CRISPR phenotypes, and upstream kinase predictions from (C). **G)** Workflow for studying the contribution of PPM1G and KHSRP phosphosites to cell survival under DNA damage. Constitutive knockdown of PPM1G or KHSRP was produced in hTERT RPE-1 dCas9-KRAB p53^−/-^ cells by CRISPRi. Cells were then nucleofected with wildtype or phosphomutant proteins for 48h time, following exposure to seven genotoxic agents, after which cell viability was measured. **H-I)** Viability of cells transfected with the indicated protein constructs, normalised to wildtype viability. Constructs were compared to wildtype within each treatment by two-way ANOVA followed by Šídák’s post-hoc tests (n = 3). *0.01 < *p* < 0.05, **0.001 < *p* < 0.01, \*\*\**p* < 0.001. nc: Negative control.

Beyond NBN and RAD51C, we predicted many high-confidence regulatory phosphosites within canonical DNA repair-linked pathways, including homologous recombination, non-homologous end joining, and DNA replication (Fig. 5b). This included proteins with no previously annotated DNA repair-regulating phosphosites, such as RAD51AP1 and TOPBP1 (Fig. 5b). We next predicted the regulation of these phosphosites by the master DNA repair kinases ATM, DNA-PK, and ATR^55^ by combining kinase motifs with phosphoproteomics of selective inhibitors^28,45,58,59^, achieving high accuracy in predicting annotated substrates of ATM (precision = 0.43, recall = 0.64) and DNA-PK (precision = 0.20, recall = 0.43), and moderate accuracy for ATR (precision = 0.24, recall = 0.23) (Fig. 5c, Fig. S5a-f). This revealed expected phosphorylation of homologous recombination machinery by ATM, as well as unexpected crosstalk from the non-homologous end-joining regulator DNA-PK to the homologous recombination proteins RAD51AP1 and NBN (Fig. 5b). Supporting this finding, recent work has demonstrated that DNA-PK can interact with the NBN-containing complex MRN^60^.

In addition to canonical DNA repair pathways, almost 40% of our high-confidence DNA repair-regulating phosphosites were in non-canonical proteins that lacked GO annotation for “DNA repair” (Fig. 5d). Encouragingly, more than half of these phosphosites were in proteins that affected viability in gene-level CRISPR screens performed under DNA damage or related conditions^61^, suggesting many may be important to the successful resolution of DNA damage (Fig. 5e). These phosphosites are typically regulated under multiple DNA damage conditions (Table S7), and many reside within the ubiquitin-proteasome system, the RNA processing systems, and splicing factors, which have been implicated in the broader DNA damage response^62–65^ (Fig. 5f). Among these predictions were UBE3A S217 and PSMD4 S266, which lack “DNA repair” annotation in PhosphositePlus but have recently been shown to mediate DNA repair through proteasome regulation^56,66,67^, further underscoring the ability of our workflow to recover biologically relevant phosphosites.

### Phosphosites on PPM1G and KHSRP regulate cell survival under DNA damage

Finally, we sought to experimentally test our predictions of non-canonical phosphosites regulating DNA repair. The protein phosphatase PPM1G and the splicing regulator KHSRP displayed multiple DNA damage CRISPR phenotypes and each contained two phosphosites predicted to regulate DNA repair (PPM1G S183, PPM1G S201, KHSRP S125, and KHSRP S274, Fig. 5f). Furthermore, two of these phosphosites have been shown to mediate secondary biological responses to DNA damage - PPM1G S183 promoted NF-kB-mediated transcription and KHSRP S274 induced miRNA biogenesis^68,69^ - however it is unclear whether these sites are also essential for cells to correctly resolve and survive DNA damage. These factors led us to examine PPM1G S183, PPM1G S201, KHSRP S125, and KHSRP S274 by site-specific mutagenesis (Fig. 5g).

CRISPRi-mediated constitutive knockdown of PPM1G and KHSRP markedly reduced the survival of hTERT RPE-1 dCas9-KRAB p53^−/-^ cells during treatment with five and seven genotoxic agents respectively (Fig. 5h-i, S6a-b, see Methods). Re-expression of wildtype PPM1G or KSHRP for 48h prior to treatment rescued viability, confirming the involvement of these proteins in the DNA damage response. We re-expressed phosphomutants for the four predicted DNA repair sites in their relevant knockdown backgrounds, and found that PPM1G S201A and KHSRP1 S125A failed to restore wildtype viability levels with multiple genotoxic agents, whereas PPM1G S183A enhanced viability beyond wildtype levels, and KHSRP S274A had no effect on viability. Immunoblotting confirmed that PPM1G S201A and KHSRP S125A were not expressed at lower levels than wildtype nor was PPM1G S183A expressed at higher levels, implying that the observed phenotypic differences were not caused by changes in protein expression (Fig. S6c-d). Together, these results suggest that phosphorylation of PPM1G S201 and KHSRP1 S125 promotes survival under several types of DNA damage, whereas phosphorylation of PPM1G S183 inhibits survival.

We next assessed the specificity of our predictions by examining two phosphosites that we did not predict to regulate DNA repair - KHSRP T143 and PPM1G S215 - since they had a low phosphosite functional score and were not quantified in DNA damage phosphoproteomics experiments (Table S8). Re-expression of KHSRP T143A restored viability to a similar extent as re-expression of wild-type KHSRP, however PPM1G S215A failed to restore viability compared to wild-type PPM1G, despite not showing decreased protein abundance as measured by immunoblotting (Fig. 5h-i, S6a-d). For all KHSRP site mutants we observed an unexpected higher molecular-weight band in immunoblotting, however its expression was also equivalent across mutants and is hence unlikely to explain their distinct effects on cell viability (Fig. S6c-d). Together, this suggests that phosphorylation of PPM1G S215, but not KHSRP T143, regulates cell survival following DNA damage. It is also possible that S215 is a functional residue mediating PPM1G’s involvement in DNA repair through a non-phosphorylation mechanism, which was disrupted in the S215A mutant.

Overall, our experiments confirmed four of six tested phosphosite predictions, comprising three of four predictions of sites regulating DNA repair and one of two predictions of uninvolved sites. This corroborates the accuracy of our computational approach, implying that further mechanistic discoveries await within the set of phosphosites we have implicated in the DNA damage response.

## Discussion

Since mass spectrometry can now routinely measure tens of thousands of phosphosites, the bottleneck in protein phosphorylation research has shifted from discovery to functional characterisation. To tackle the problem of predicting the cellular functions of phosphosites, we developed predictors of phosphosite-regulation for six major cellular processes, and experimentally validated DNA repair predictions for phosphosites on KHSRP and PPM1G. We anticipate that our predictions will aid interpretation of phosphoproteomics experiments and will provide a springboard for deeper investigations into phosphosite function.

A major strength of our predictions was their interpretable features, including phosphosite regulation across diverse, biologically relevant phosphoproteomics perturbations. Furthermore, our phosphoproteomics atlas was crucial for improving model accuracy beyond that afforded just by protein-level functional annotation, underscoring the utility of reusing omics experiments in large-scale compilations. However, our atlas was sparse with each perturbation quantifying on average only 8,300 of the 189,576 total phosphosites, and this limited the scale at which we could train and apply our predictors. This likely reflects biological factors - only a subset of possible sites will be phosphorylated in a given context - as well as technical limitations of mass spectrometry, especially since many datasets in the atlas were acquired with older instruments and methodologies. Hence, to gain more comprehensive and richer insights, we will need concerted efforts to produce large-scale phosphoproteomics datasets, leveraging recent advances in mass spectrometry instrumentation and phosphoproteomics workflows that increase quantitative depth and reproducibility^70,71^. Furthermore, just as it is now a standard publishing requirement to share raw data in repositories such as PRIDE^72^, the reusability of (phospho)proteomics data will be further enhanced if authors are required to deposit analyzed data in a standardized format and with rich meta-information.

Reasoning that protein kinases orchestrate specific cellular pathways, we incorporated kinase sequence specificities as a feature in our models. However, with the exception of DNA damage-sensing kinases, these could only weakly predict phosphosite cellular function. This is not entirely surprising, given that the sequence specificities of many functionally divergent kinases are highly degenerate^16^. Still, it is intriguing that sequence preferences have evolved as a determinant of functional specificity for only one small set of kinases. Presumably, functional specificity is imparted by other mechanisms of kinase-substrate specificity, such as distal docking motifs, protein interaction partners and subcellular localisation^73^. Recent studies in yeast and mice also demonstrated that in spite of their highly similar sequence motifs, different DNA damage kinases can target distinct substrates using non-canonical, kinase-specific motifs, and this principle may apply to other kinases with similar sequence preferences^74,75^. As prediction and high-throughput discovery of kinase substrates continue to improve^43,76–78^, kinase-substrate relationships could also be incorporated as a feature to improve predictions of phosphosite cellular function.

It is likely that the accuracy of our models could be improved by incorporating additional features as they become available. For example, since cellular processes are often spatially compartmentalized, the subcellular location where a phosphosite is phosphorylated could indicate its function. Phosphoproteomics workflows are beginning to map phosphosite subcellular localisation at scale^79,80^. Furthermore, protein phosphorylation often modulates protein-protein interactions, and the functional identity of interaction partners regulated by a phosphosite could illuminate the site’s cellular function. Protein language models have proven useful in predicting residue-level properties such as signal peptides^81^ and kinase-phosphosite recognition^82^, however their utility for our task may be limited, given language model-based predictions of protein cellular function are generally worse than molecular function^83^. Finally, high-throughput experimental manipulation of phosphosites coupled to rich phenotypic readouts, such as cellular fitness across diverse stresses^8^, will help validate and further train functional predictors.

We predicted some cellular processes with high accuracy (e.g. DNA repair, cytoskeletal reorganisation), while others were hardly better than random (e.g. cell growth, differentiation). This may be because the latter processes are broad and encompass many distinctly regulated sub-processes. Higher accuracy may be obtained on more fine-grained and potentially more functionally coherent phosphosite annotations, which would also expand our predictions from six general processes to a richer functional map. It is also important to know whether a predicted relationship is activating or inhibitory; while such directionality is included in PhosphositePlus annotations, it is absent for many phosphosites and conflicted for others (Fig. S7a), hence we decided to exclude it from this study. Finally, it is worth noting that existing annotations, both at the site level and protein function level, are likely biased to highly studied proteins^84^. However, our predictions do not seem to be confined to such proteins, given that many high-confidence predictions were on understudied proteins lacking the corresponding GO-term annotation. In all, in this study we have demonstrated that existing annotations can be used to accurately predict cellular functions of phosphosites. As annotation databases expand in depth and detail, and as more experimental data becomes available, such predictions will only increase in their scope and accuracy.

## Acknowledgements

J.E.C. is supported by the NOMIS Foundation, the Lotte and Adolf Hotz-Sprenger Stiftung, the Swiss National Science Foundation (project grants 310030_188858, 320030-227979, and 310030_201160), and the European Research Council (ERC) under the European Union’s Horizon 2020 research and innovation program (grant agreement No 855741, DDREAMM). D.G. is supported by a Helen Hay Whitney Foundation Postdoctoral Fellowship (ID F1290).

## Author contributions

Conceptualization: P.B., J.vG., Data curation: J.vG., Formal analysis: J.vG., Funding acquisition: P.B., J.E.C., Investigation: J.vG., D.G., I.S., J.J., K.O., Supervision: P.B., J.E.C., Visualization: J.vG. Writing – original draft: J.vG., Writing – review & editing: All authors

## Data availability

The phosphoproteomics compilation has been deposited to BioStudies with the identifier S-BSST2989, and is accessible at www.ebi.ac.uk/biostudies/studies/S-BSST2989.

## Code availability

All code used to analyse data and produce figures has been deposited on GitHub at the repository github.com/JulianvanGerwen/PhosphositeCellularFunctions.

## Methods

### Analysis

#### Computational analysis

Unless otherwise studied, analysis was performed using R (version 4.3.1) in the RStudio environment (version RStudio 2024.09.1+394).

#### PhosphositePlus annotations

Regulatory phosphosite annotations were downloaded from PhosphositePlus (17 July 2023). Cellular process annotations were extracted from the “ON_PROCESS” column and the directionality of regulation was removed. Analysis was performed on all human phosphosites annotated to regulate at least one cellular process. In figures and text the following abbreviations were used instead of the original PhosphositePlus labels: “cell cycle” instead of “cell cycle regulation”; “signaling” instead of “signaling pathway regulation”; “cytoskeleton” instead of “cytoskeletal reorganization”; and “differentiation” instead of “cell differentiation”.

#### ROC-based prediction of cellular function

In general, ROC curve analysis was used to evaluate the ability of features to predict the cellular processes regulated by phosphosites. For a given process, the positive class was defined as all phosphosites annotated to regulate the process, and the negative class was defined as all phosphosites annotated to regulate at least one other process. Area under the ROC curve (ROC AUC) was calculated using roc_auc() from the R package yardstick (version 1.2.0).

#### Gene ontology (GO) analysis

Gene ontology (GO) biological process terms were extracted using gene symbols and the GOfuncR package (version 1.20.0). Annotations were propagated up the GO hierarchy, such that genes were annotated to all GO terms hierarchically above their original annotated terms. Each gene’s GO terms were assigned to its phosphosites. For each cellular process, GO term enrichment was performed by one-side Fisher’s exact test on process-regulating phosphosites, with the background as all phosphosites annotated to any process. Only GO terms containing at least three process-regulating phosphosites were tested. P-values were adjusted for each process by the Benjamini-Hochberg procedure.

#### Kinase sequence specificity analysis

Position weight matrices (PWMs) of Ser/Thr kinase sequence specificities were extracted from Johnson et al.^16^ (sheet “ser_thr_all_norm_scaled_matrice” in Table S2). Phosphoacceptor preferences were added to matrices as described in Johnson et al. For a given phosphosite, kinase motif scores were calculated by multiplying the weight values for each surrounding amino acid in its position relative to the phosphoacceptor, followed by log-transformation.

### Phosphoproteomics atlas

#### PubMed search

A PubMed search was performed to identify papers containing human phosphoproteomics data (12 Dec 2023). An advanced search was used ((((phosphoproteomics[Title/Abstract]) OR (phospho-proteomics[Title/Abstract])) OR (phosphoproteome[Title/Abstract])) OR (phospho-proteome[Title/Abstract])) NOT (Review[Publication Type]). Additional filters were used: “Human” for species and publication date > 2015. From the 1,234 resulting studies, 27 were selected because they contained many different conditions, or conditions deemed relevant for predicting phosphosite cellular function. Studies present in the Ochoa 2016 or qPTM compilations were excluded^18,19^.

#### Data extraction

Phosphoproteomics data were extracted either by downloading supplementary tables when available, or downloading searched spectral data from the PRIDE repository^72^. When authors had already calculated fold change values between perturbations and control conditions, these were used. Otherwise, phosphopeptide log2-intensity values were normalized by subtracting sample medians, phosphopeptide means were calculated across samples in the same condition, and fold change values were calculated by subtracting means.

#### Combining data

To combine datasets, phosphopeptides were mapped into the same uniprot human proteome (UP000005640_2024_01_24) in a multi-step process.

1. Uniprot identifier selection: All datasets provided a uniprot identifier and phosphopeptide position in the corresponding protein sequence. When multiple alternative uniprot identifiers were provided for the same phosphopeptide, those in the reviewed human proteome were selected.
2. Phosphoacceptor amino acid comparison: Amino acids corresponding to reported phosphosite positions were extracted and compared to the phosphoacceptor amino acid provided by authors.
3. Peptide sequence/surrounding sequence mapping: When the mapped and provided amino acids did not agree, or when no reviewed uniprot identifier could be selected in step 1, phosphopeptide sequences were directly mapped into the human proteome. When phosphopeptide sequence or phosphosite surrounding sequences were not provided, the corresponding phosphopeptides were removed. When a phosphopeptide matched multiple positions in the proteome, the first uniprot identifier was selected, prioritising reviewed identifiers were available. Phosphopeptides that could not be mapped into the human proteome were removed.

#### Phosphopeptide processing and data filtering

Each phosphopeptide was identified by its uniprot identifier, phosphosite positions, and multiplicity of phosphorylation. When multiple phosphopeptides had the same values for these identifiers, quantitative information was collapsed by taking the mean of log2 fold-change values. Log2 fold change values representing technical or biological replicates within the same study were averaged. Analysis was restricted to mono-phosphorylated peptides for ease of interpretation, and because multiply phosphorylated peptides were handled differently in different studies. Conditions with 2,000 or fewer quantified fold change values were removed. To mitigate false positive identifications, phosphosites quantified in only one condition were removed.

#### Processing qPTM data

Phosphoproteomics studies performed in human samples were extracted from the qPTM compilation^18^. Phosphopeptides with no provided sequence position were removed. Log2 fold change values representing technical or biological replicates within the same study were averaged. Values from phosphopeptides with the same uniprot identifier, phosphosite positions, and multiplicity of phosphorylation were averaged. Phosphopeptides with uniprot identifiers not present in the uniprot human proteome were removed. Phosphosite sequence windows were extracted from the uniprot human proteome and removed when they did not match those provided in qPTM. Data was filtered for mono-phosphorylated peptides, conditions with more than 2,000 quantified fold change values, and phosphosites quantified in more than one condition.

#### Processing Ochoa 2016 data

ENSP identifiers were mapped to uniprot identifiers using Ensembl BioMart. Phosphopeptide sequences were mapped into corresponding uniprot sequences to extract phosphosite positions. Values from phosphopeptides with the same uniprot identifier, phosphosite positions, and multiplicity of phosphorylation were averaged. Data was filtered for mono-phosphorylated peptides, conditions with more than 2,000 quantified fold change values, and phosphosites quantified in more than one condition.

#### Combining compilations

The pubmed search-derived phosphoproteomics compilation was combined with qPTM and Ochoa 2016. When the same study was present in qPTM and Ochoa 2016, the values from qPTM were used.

### Model training and testing

#### Train-test splits

Models were trained and tested on 10 random iterations of five-fold cross-validation, whereby PhosphositePlus regulatory sites were split into five folds, each fold was retained once as a test set, and models were trained on the remaining four folds. To prevent data leakage from feature selection of protein-level GO terms on training data, phosphosites belonging to the same protein were grouped and folds were constructed at the protein level, with proteins assigned to cellular processes when they contained at least one process-regulating phosphosite. To maintain the relative proportions of different cellular processes in training and test data, stratified five-fold splits were performed using the iterative.stratification.partitions() function from the R package mldr.datasets (version 0.4.2).

#### Feature selection

Within training sets, features were selected for each cellular process as follows:

- GO biological process enrichment was performed as described above. The most significantly enriched GO term (smallest adjusted p-value) was selected. When multiple GO terms were tied, the term containing more process-regulating phosphosites was selected.
- Ser/Thr phosphosites were scored with each Ser/Thr kinase PWM as described above. ROC AUCs were calculated for predicting process-regulating phosphosites with each PWM. The kinase PWM with the highest AUC was selected.
- Phosphoproteomics log2 fold-change values were converted into absolute values. ROC AUCs were calculated for predicting process-regulating phosphosites with absolute log2 fold change values from each condition, among quantified Ser/Thr phosphosites. Conditions with fewer than 26 phosphosites quantified in the positive or negative class were removed. The 20 conditions with the highest AUC values were selected. Conditions with AUC < 0.61 were excluded. Correlated conditions were removed using the findCorrelation() function from the R package caret (version 6.0-94, filtering r > 0.5).

#### Model training

Training and test data were subsetted for Ser/Thr phosphosites quantified in at least three selected phosphoproteomics conditions. Missing values in phosphoproteomics data were imputed using the mean absolute log2 fold change in each condition. All features were centred and scaled using the preProcess() function from the R package caret (version 6.0-94). Binary predictors were trained using class weights inversely proportional to class frequency to account for class imbalance. Model hyperparameters were selected based on the highest ROC AUC in the default grid search provided by caret, using five-fold cross validation on the training set repeated 10 times. The train() function from caret was used to train multiple architectures with the specified parameters: logistic regression (method = “glm”, family = “binomial”); gradient-boosting machine (method = “gbm”); and random forest (method = “ranger”). Model performance was evaluated by ROC AUC values on test sets.

#### Predicting

Logistic regression models from each train-test split were applied to all Ser/Thr phosphosites quantified in at least three selected phosphoproteomics conditions, with Ochoa 2020 functional score > 0.5^3^. Final predictions for each process were obtained by taking the median across all 50 models from train-test splits, only considering phosphosites predicted by at least 80% of models. To set a cutoff for high-confidence phosphosites, model false-positive rates (FPR) were estimated on each test set across a range of score cutoffs, and the lowest score cutoff that gave a median FPR less than or equal to 5% was selected.

#### Feature weight

Feature weight for each cellular process-predictor was calculated by summing the absolute values of logistic regression coefficients across all 50 models from train-test splits.

### CPTAC multiomics data

#### Data extraction and processing

CPTAC data were extracted using the cptac python package (cptac version 1.5.14, python version 3.12.4). The following pipelines were used: phosphoproteomics - bcm; proteomics - bcm; transcriptomics - bcm; whole-exome sequencing - harmonized. Non-tumour samples were removed. For phosphoproteomics, proteomics, and transcriptomics, data were transformed into log2 fold changes by subtracting the median within each tumour type for each quantified phosphosite/protein/transcript. Somatic mutation burden was calculated by counting the number of mutations called for each tumour in whole-exome sequencing data.

#### Transcription factor and DNA repair activity estimation

Transcription factor targets were extracted from the DoRothEa database (R package dorothea^85^ version 1.12.0, confidence scores A-C). Transcription factor activity was estimated using the VIPER algorithm (viper() function from the R package viper^86^ version 1.34.0) with minsize = 5, adaptive.size = FALSE, and eset.filter = F. DNA repair signaling activity was estimated from DNA repair-regulating phosphosites using the geneSetTest() function from the R package limma (version 3.56.2) with alternative = “either”, type = “t”, and ranks.only = TRUE. Phosphosite log2 fold changes were supplied as statistics. The resulting p-values were log10-transformed and signed by the mean fold-change of DNA repair phosphosites, such that up-regulated phosphosites resulted in a positive value and down-regulated phosphosites resulted in a negative value.

### DNA repair analysis

#### ClinVar data

Variants from ClinVar (16/04/2024) were mapped to the proteome using ProtVar^87^, and filtered to keep missense variants with at least one review star. Only missense mutations were considered, and mutations that were not linked to specific diseases (i.e. the CLNDN value was “not_provided” or “not_specified”) were removed.

#### Curating canonical DNA repair proteins

Canonical DNA repair proteins were curated using genes annotated under the GO terms “double-strand break repair via homologous recombination” and “double-strand break repair via nonhomologous end joining”, combined with a literature search. The DNA replication genes MCM3 and MCM6 were added.

#### DNA repair kinase substrate prediction

Substrates of ATM, DNA-PK, and ATR were predicted using sequence specificity scores calculated as described above and phosphoproteomics data of selective kinase inhibitor treatment under DNA damage conditions (ATM: Schlam-Babayov et al.; DNA-PK: Justice et al.; ATR: Schlam-Babayov et al., Wagner et al., Salovska et al.^28,45,58,59^). To account for data sparseness, the most negative log2 inhibitor/control fold-change value was calculated. Phosphosites with a motif score > 1 and log2 inhibitor/control fold-change < -1 were classified as kinase substrates.

### DNA damage experiments

#### Site directed mutagenesis

Point mutation of phosphosites was performed using mutagenic primers that encoded the mutation of interest (listed below). PCR amplification of parental plasmid (KHSRP: Addgene 23001; PPM1G was cloned using a gBlock ordered from IDT) was performed with mutagenic primers using Q5 high-fidelity DNA polymerase. Following amplification, PCR reactions were digested using Dpn1 at 37°C for 1 hour. Following SPRI bead purification, DNA was transformed into STBL3 chemically competent *E. coli* and plated onto antibiotic containing agar plates. Colonies were sequenced to confirm incorporation of point mutation.

KHSRP_T143A: TCCCCCAAGGgcgTCAATGACAG

KHSRP_S125A: GAAGCTGGCTgcgCAGGGAGACT

KHSRP_S274A: TCAGGACGGAgcgCAGAATACGA

PPM1G_S183A: GGAACCAGGGgcgCAGGGCCTCA

PPM1G_S201A: GGAAACTCCTgcgCAAGAAAATG

PPM1G_S215A: CACAGGCTTTgcgTCCAACTCGG

#### Cell Culture

The hTERT RPE-1 dCas9-KRAB p53 null and HEK293T cell lines were cultured in Dubecco’s modified Eagle’s medium/F12 (Merck) supplemented with 10% fetal bovine serum (thermofisher) and 100 units/mL of streptomycin and 100 mg/mL of penicillin (Gibco). Following transduction with lentivirus, hTERT RPE-1 dCas9-KRAB p53 null cells were selected with media containing puromycin at 15 ug/mL. Cells were routinely tested for mycoplasma.

#### Guide RNA cloning

CRISPRi gRNAs were cloned into a puromycin-resistance-containing lentiviral vector (Addgene 60955). Forward and reverse oligos for each sgRNA were annealed by first incubating at 37°C for 30 minutes with T4 polynucleotide kinase (NEB), following by incubation at 95°C for 5 minutes and then ramping down to 25°C by 1 degree per minute. Annealed sgRNAs were ligated into the *BstXI* and *Blp1* digested backbone using T4 ligase (NEB). sgRNA protospacers were selected from the Weissman CRISPRi library.

#### Lentivirus packaging

Lentiviruses were produced in HEK293T cells. In brief, HEK293T cells were transfected using polyethylenimine with the VSV-G envelope expressing plasmid (pMD2.G), packaging plasmid (psPAX2) and the cloned sgRNA-containing transfer vector. Lentiviral supernatant was collected 48 hours after transfection.

#### Complementation Assays

hTERT RPE-1 dCas9-KRAB p53 null cells transduced with either a KHSRP or PPM1G sgRNA were nucleofected with a plasmid containing a wild-type allele of the corresponding knockdown or the indicated point mutant using a Lonza Amaxa nucleofector, P3 solution, and program EA104. Cells were grown in non-selection media for 48 hours to allow for expression of the neomycin resistance cassette encoded on the nucleofected plasmid, after which nucleofected cells were selected by adding G418 to the media at a concentration of 1.25 mg/mL. Following selection, cells were seeded at 8,000 cells/well of a 96-well plate and treated with the indicated drugs as performed previously^88^ (Aphidicolin 1uM; Bleomycin 5ug/mL; Etoposide 500 nM; Hydroxyurea 1 mM; MitomycinC (MMC) 5 nM, Methyl Methanesulfonate (MMS) 0.005%) for 18 hours and then washed out. For Olaparib conditions, cells were treated chronically at 2 uM. One week following treatment, cell viability was measured using CellTiter-Glo luminescence and measured using a Victor Nivo plate reader. Due to inconsistencies with cell viability readouts in the knockdown studies of PPM1G, treatments with Aphidicolin and Etoposide were removed from the study.

#### Immunoblotting

Cells were lysed in radioimmunoprecipitation assay buffer (RIPA) (0.5M Tris-HCl, 1.5 M NaCl, 2.5% deoxycholic acid, 10% NP-40 and 10 mM EDTA), supplemented with Halt Protease Inhibitor Cocktail and Phosphatase Inhibitor Cocktail (ThermoFisher Scientific). Samples were sonicated using a Bioruptor Plus probe-based sonicator (30s ON and 30s OFF for 5 cycles) and centrifuged for 5 minutes at 21,000g. Protein concentrations were measured using a bradford assay with absorption measured on a Nivo plate reader. The samples were normalized by protein concentration, mixed with NuPAGE LDS Sample buffer and 5% β-mercaptroethanol, and boiled for 5 minutes at 95°C. The samples were loaded into a Bolt 4-12% Bis-Tris Protein Gel (ThermoFisher Scientific) and transferred to a 0.2-µm nitrocellulose membrane. Membrane blocking was performed using 5% milk diluted in Tris-buffered saline containing 0.1% Tween 20. After incubation with primary antibodies, membranes were incubated with Li-Cor near-infrared fluorescence secondary antibodies, after which they were scanned using a Li-Cor Near-InfraRed fluorescence Odyssey CLx Imaging System. All primary antibodies were used at a dilution of 1:1000 and secondaries were used at a concentration of 1:15,000. Primary antibodies used were: Rabbit anti KHSRP from Novus Biologicals (NBP1-18910), Rabbit anti PPM1G from Novus Biologicals (NB110-38867), Rabbit anti GAPDH from Cell Signaling Technology (2118S).

## Supplementary figures

**Figure S1.**
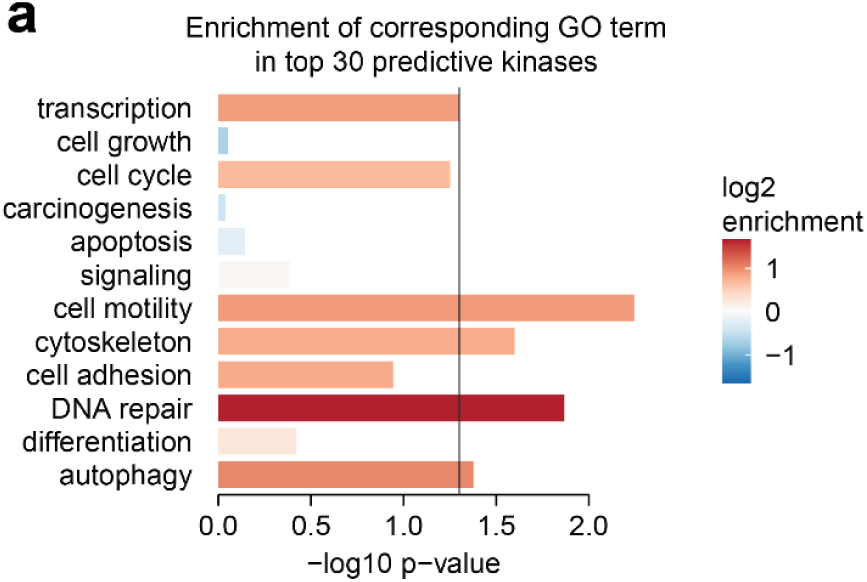
**A)** For each cellular process, the GO term most significantly enriched in process-regulating phosphosites was selected as in Fig. 1c). The 30 Ser/Thr kinase motifs with the highest ROC AUC for predicting process-regulating phosphosites were selected. One-sided Fisher’s exact tests were performed to test the enrichment of the selected GO term among these kinases, with all 303 profiled Ser/Thr kinases as the background.

**Figure S2.**
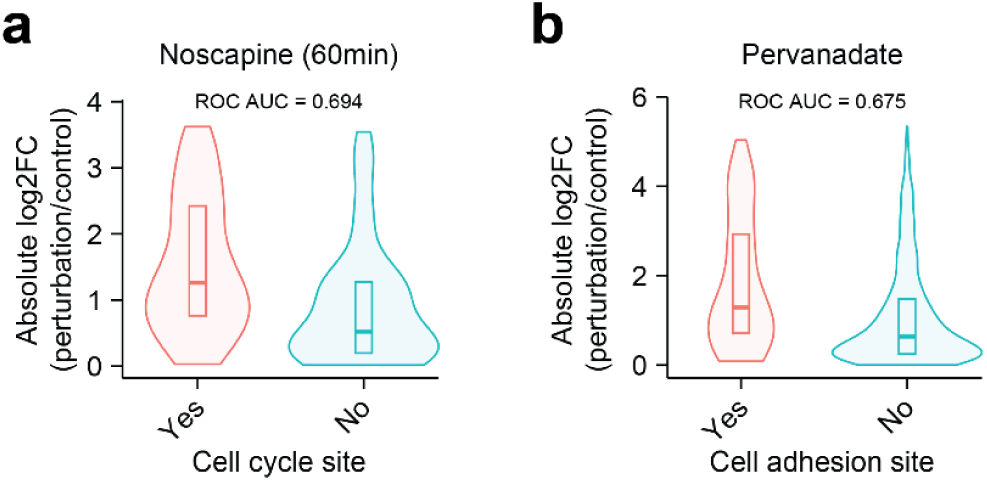
**A)** Cell cycle-regulating phosphosites are preferentially regulated under noscapine treatment. **B)** Cell adhesion-regulating phosphosites are preferentially regulated under pervanadate treatment.

**Figure S3.**
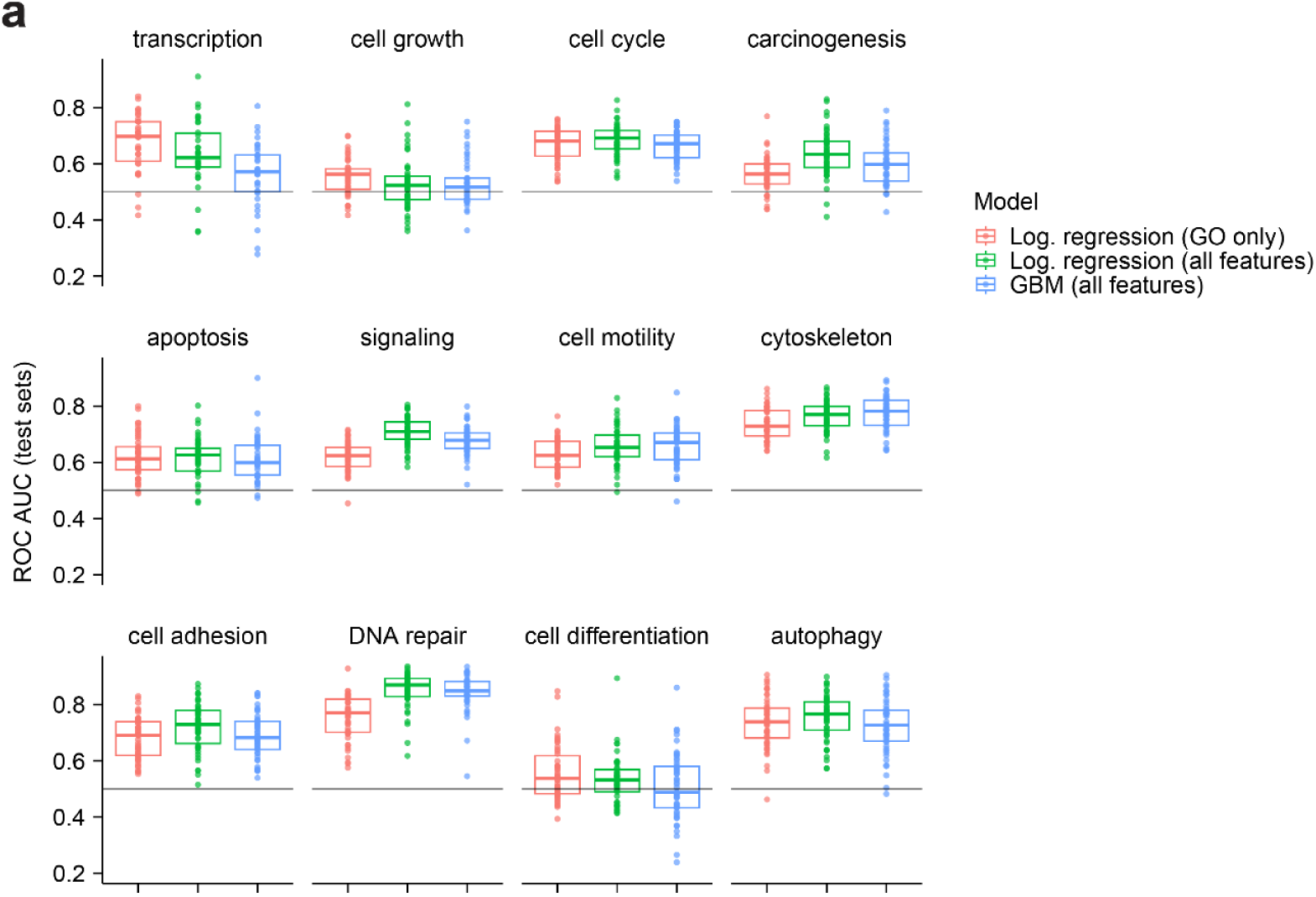
**A)** Phosphosite cellular function predictors were trained using logistic regression (“log. regression”) with GO terms, kinase motifs, and phosphoproteomics perturbations (“all features”, also shown in Fig. 3b), gradient-boosting machines with all features (“GBM”), or logistic regression only using GO terms. ROC AUC values over 50 train-test splits are shown. The same set of train and test phosphosites were used for all model types.

**Figure S4.**
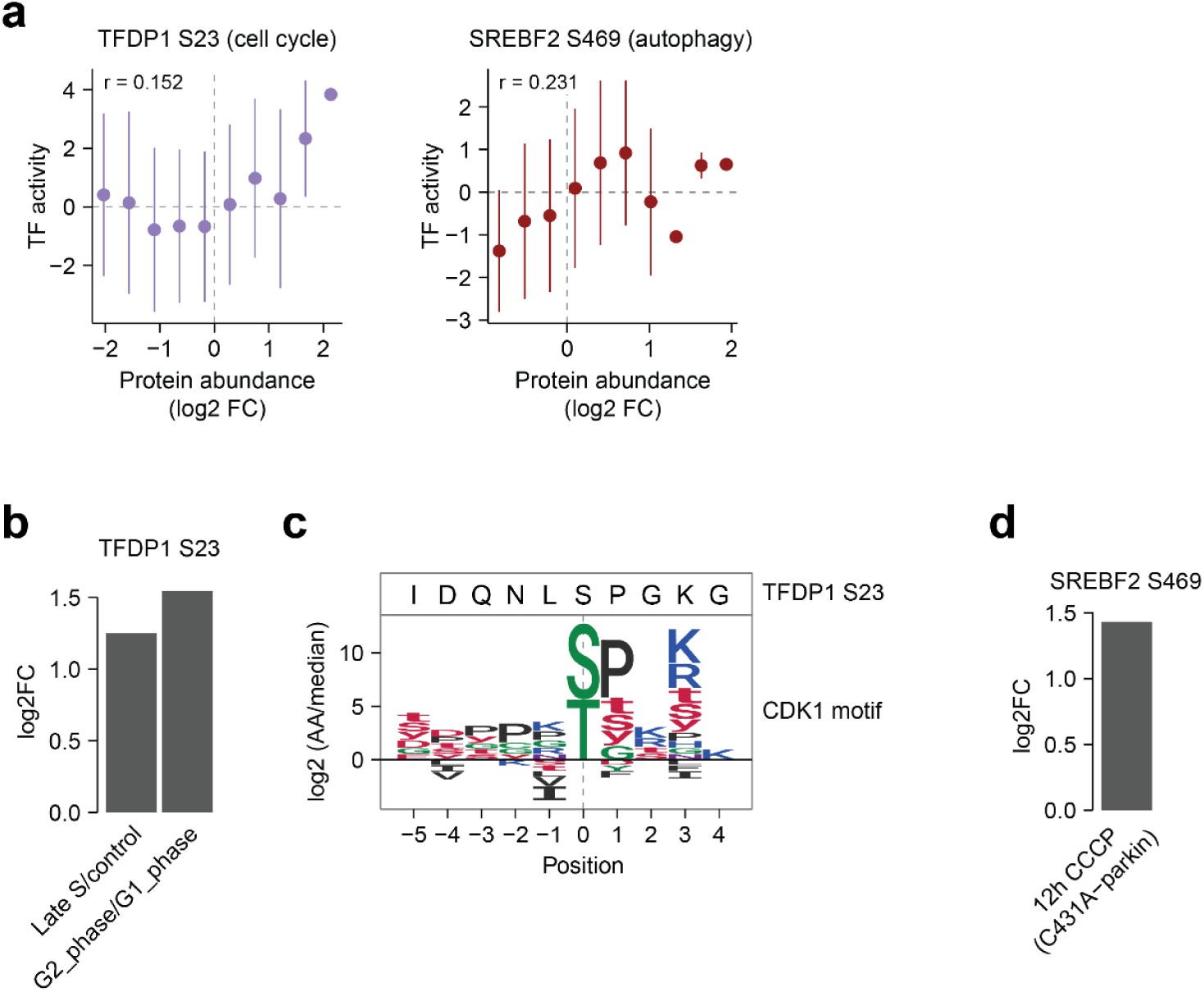
**A)** Pearson’s correlation between transcription factor activity and protein abundance for phosphosites associated with transcription factor activity. Points represent mean transcription factor activity in bins of protein abundance and error bars represent standard deviation. **B)** Regulation of TFDP1 S23 in selected phosphoproteomics conditions. **C)** The surrounding sequence of TFDP1 S23 aligned with the CDK1 sequence motif^16^. **D)** Regulation of SREBF2 S469 in a selected phosphoproteomics condition. LogFC is calculated relative to a 0-hour control on a background of C431A-parkin overexpression.

**Figure S5.**
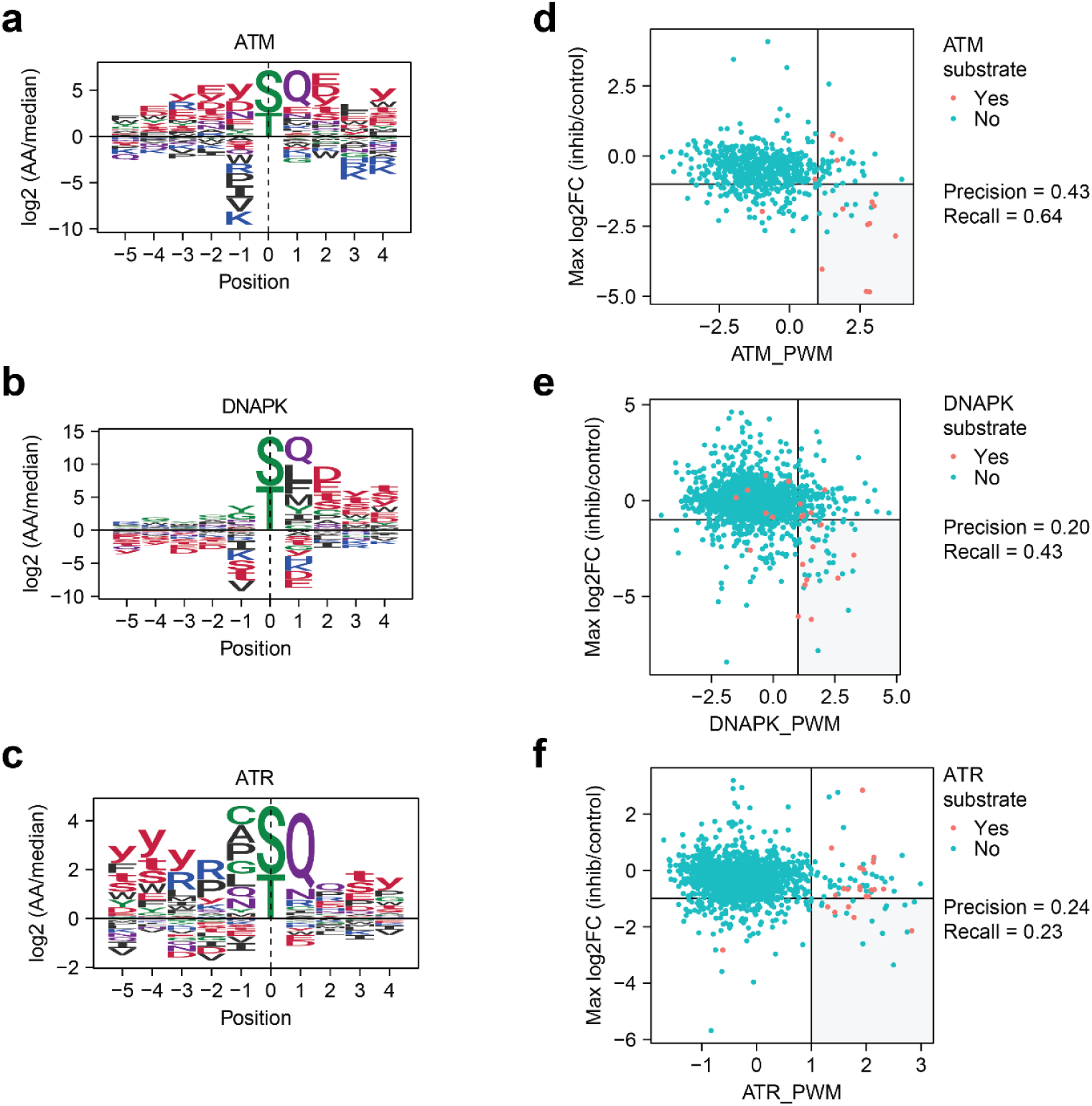
**A-C)** The sequence motifs for ATM, DNAPK, and ATR^16^. **D)** The ATM sequence motif and ATM inhibitor phosphoproteomics data were used to predict ATM substrates. The x-axis shows ATM motif scores and the y-axis shows the most negative log2 fold change across different ATM inhibitor treatments (See methods). All phosphosites annotated to at least one upstream kinase in PhosphositePlus are shown, and ATM substrates are colored red. Phosphosites with motif score > 1 and inhibitor log2 fold change < -1 were classified as ATM substrates, with precision and recall shown. **E-F)** As in D), but for DNA-PK and ATR.

**Figure S6.**
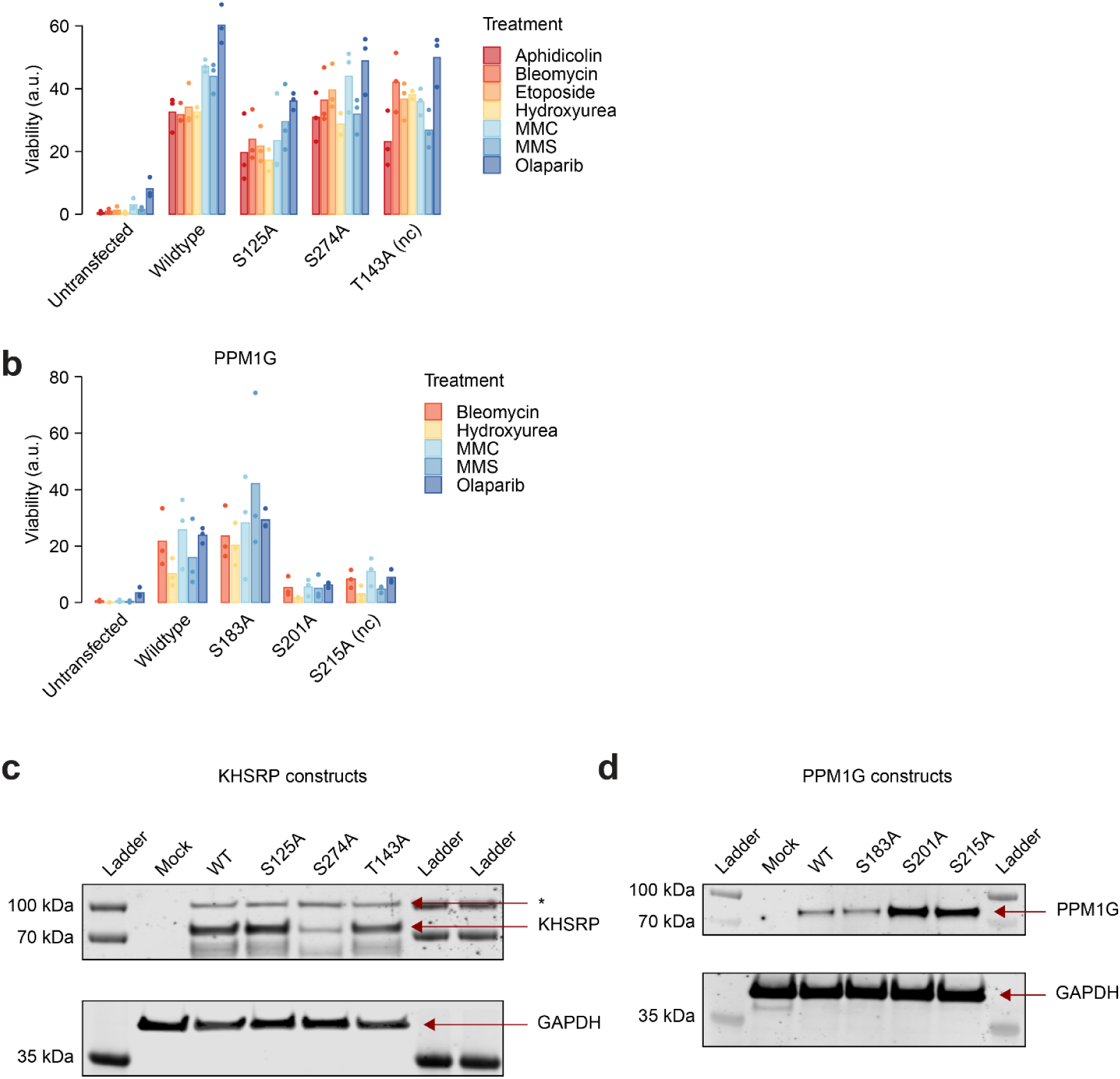
**A-B)** Un-normalised viability results, related to Fig. 5h-i. Constitutive knockdown of PPM1G or KHSRP was produced in hTERT RPE-1 dCas9-KRAB p53^−/-^ cells by CRISPRi. Cells were then nucleofected with wildtype or phosphomutant proteins for 48h time, following exposure to seven genotoxic agents, after which cell viability was measured. **C-D)** Immunoblotting of KHSRP, PPM1G, and GAPDH (loading control) from cells nucleofected with the indicated constructs or without nucleofection (Mock). * indicates an unexplained higher-molecular weight band above KHSRP.

**Figure S7.**
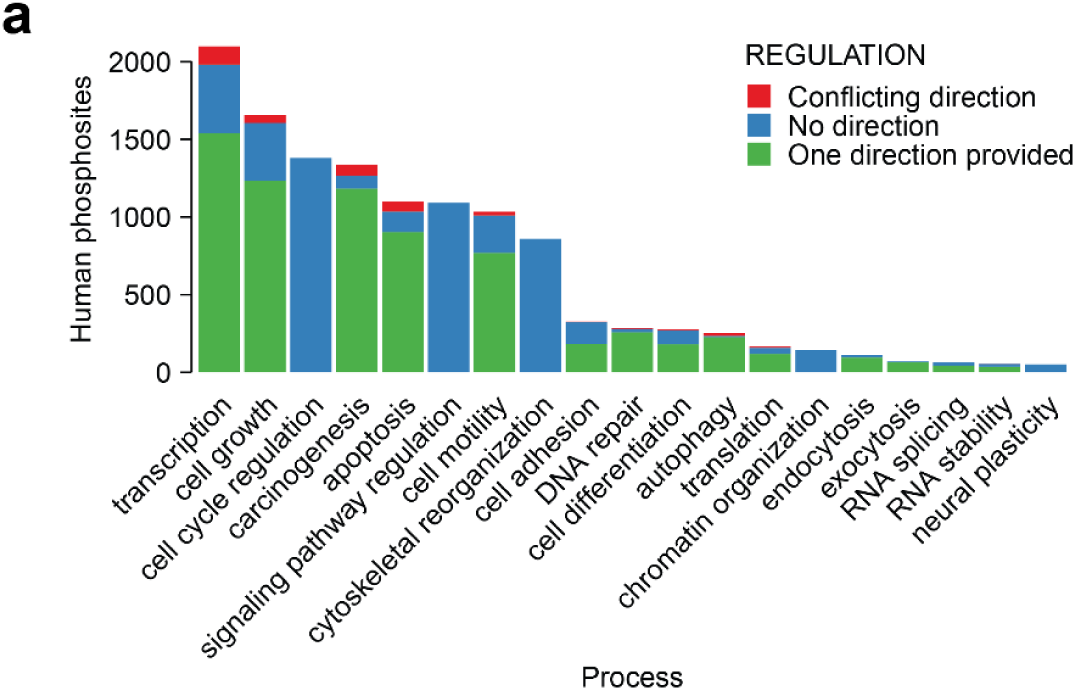
**A)** Human phosphosites annotated to regulate each cellular process in PhosphositePlus. Phosphosites are classified based on whether one direction of regulation was provided (“induced” or “inhibited”), no direction was provided (“altered”), or two directions were provided (“induced” and “inhibited”).

## Supplementary tables

**Table S1**: Annotated regulatory phosphosites from PhosphositePlus All human phosphosites annotated to regulate one or more cellular processes. The processes regulated by each phosphosite are indicated with the value “1” in the corresponding cell.

**Table S2**: Biological Process terms enriched in process-regulating phosphosites For each cellular process annotated in PhosphositePlus, GO Biological Process enrichment was performed on process-regulating phosphosites. The top 20 most significantly enriched GO terms are shown for each process.

**Table S3**: Condition metadata for phosphoproteomics compilation Metadata for all conditions in the phosphoproteomics compilation.

**Table S4**: Phosphosite cellular function predictions All phosphosite cellular function prediction scores. The general phosphosite functional score from Ochoa 2020 is also shown.

**Table S5**: Features selected in models Features selected by phosphosite function models. The number of logistic regression models in which each feature was selected is shown (maximum 50), as well as the overall feature weight (sum of absolute coefficients across models).

**Table S6**: DNA repair phosphosite enrichment in CLINVAR diseases Enrichment was performed to link predicted DNA repair-regulating phosphosites to CLINVAR genetic diseases. First, phosphosites harbouring disease-linked missense mutations were collected. Then, rank-based enrichment (one-sided geneSetTest, alternative = “greater”. R package limma version 3.56.2) was performed to identify diseases with phosphosites featuring high DNA repair prediction scores.

**Table S7**: Regulation of DNA repair predictions under DNA damage High confidence DNA repair predictions on genes with DNA damage CRISPR phenotypes and without DNA damage GO term annotation were collected. Their regulation across 14 DNA damage phosphoproteomics conditions is shown (log2 fold changes).

**Table S8**: Phosphosites experimentally validated for involvement in DNA repair Information for phosphosites on PPM1G and KHSRP that were experimentally validated for involvement in DNA repair. Experiments were performed on four phosphosites predicted to regulate DNA repair and two phosphosites not predicted to regulate DNA repair. The Ochoa 2020 phosphosite functional score is shown, as well as phosphosite regulation across 14 DNA damage phosphoproteomics conditions (log2 fold changes).

